# Change in prey genotype frequency rescues predator from extinction

**DOI:** 10.1101/2022.04.19.487766

**Authors:** Ruben Joseph Hermann, Lutz Becks

**Affiliations:** Limnological Institute University Konstanz, Aquatic Ecology and Evolution Group. Konstanz, Germany

**Keywords:** predator-prey, indirect evolutionary rescue, rotifer, *Chlamydomonas*, frequency dependency, eco-evolutionary dynamics

## Abstract

Indirect evolutionary rescue (IER) is a mechanism where a non-evolving population is saved from extinction in an otherwise lethal environment by evolution in an interacting population. This process has been described in a predator-prey model, where extinction of the predator is prevented by a shift in the frequency of defended towards undefended prey when reduced predator densities lower selection for defended prey. We test here how increased mortality and the initial frequencies of the prey types affect IER. Combining the analysis of model simulations and experiments with rotifers feeding on an algal population we show IER in the presence of increased predator mortality. We found that IER was dependent on the ability of the prey population to evolve as well as on the frequency of the defended prey. High initial frequencies of defended prey resulted in predator extinction despite the possibility for prey evolution, as the increase in undefended prey was delayed too much to allow predator rescue. This frequency dependency for IER was more pronounced for higher predator mortalities. Our findings can help informing the development of conservation and management strategies that consider evolutionary responses in communities to environmental changes.

## 1. Introduction

Evolutionary rescue (ER) describes the process where a population evolves to avert extinction in an otherwise lethal environment [1,2] through the increase of an adapted genotype with a positive net growth rate. The adapted genotype may be present at low frequency [3], introduced via migration [4,5], the result of genetic mixing [6] or evolve de-novo [7,8]. As populations interact with other populations, it is, however, possible that changes in species interactions and indirect ecological and evolutionary processes affect the demography of a population [9,10] and thus the probability of rescue. Theoretical studies have started to integrate the effect of multispecies interaction into research on ER [11–16]. There are however only few studies exploring ER in communities with multiple species [17,18] and the conditions under which populations can be rescued through evolutionary change in communities are still poorly explored [19].

Predator-prey interactions are important driver for population dynamics and intraspecific diversity [9,20] and have been shown to affect species persistence of prey and/or predator through direct and indirect processes [10,21]. Indirect evolutionary rescue (IER), where a population is rescued from extinction through evolution in an interacting population has been suggested as a mechanism promoting predator persistence in an otherwise lethal environment [12]. In their model, Yamamichi and Miner [12] investigated whether heritable trait variation in the prey population that determines the interaction with the predator (i.e., defended and undefended prey) and competition between prey clones can rescue the predator from an environmental change that results in increased mortality for the predator. They found that the predator goes extinct with increasing mortality when neither prey nor predator can adapt. However, the predator can survive and gain positive population growth in the presence of increased mortality when the prey population can evolve in response to changes in the predator density. This is because decreased predator density disfavors defended prey and favors more competitive undefended prey, which in return leads to an increase in predator growth and eventually rescue of the predator population. IER in a predator-prey system leads thus to a u-shaped temporal change in the predator density following the environmental change similar to ER [1,2], but the change depends on a shift in the frequencies of the defended and undefended prey for IER.

A key assumption for the IER in predator-prey systems with at least two prey clones is that changes in the frequencies of the prey clones can prevent extinction of the predator when increased predator mortality reduces selection for defended prey and leads to an increase in the fraction of undefended prey. As fitness and thus the persistence of the predator depends on the change in the prey population, IER is expected to be a function of the frequency of the defended prey and density of total prey at the time when the environment becomes lethal. For example, when the frequency of the undefended prey is low to begin with, the predator might go extinct before the undefended prey reaches high enough densities. Or when the frequency and density of the undefended prey is initially high, the predator might survive without evolutionary change in the prey population. We tested here the frequency dependency of IER by combining the analyses of model simulations and experiments. First we identified combinations of mortality rates of the predator and frequencies of defended prey under which IER occurs by adapting a predator-prey model from Becks *et al*. [20]. In a second step, we tested for IER in experiments using a freshwater predator-prey system consisting of the rotifer *Brachionus calyciflorus* feeding on the green algae

### Chlamydomonas reinhardtii

To manipulate the presence/absence of evolution in the prey population as well as the frequency of defended prey, we used two different isogenic lines of *C. reinhardtii*. The clones differed in their defense trait against the rotifer, where the higher defense correlated with a lower growth rate (Fig. S1) [22]. We used Sodium Chloride (NaCl) as a stressor because it increases the rotifer mortality but has no significant effect on algal growth at the concentrations considered here (Figs. S2, S3). By varying NaCl concentrations, we tested a range of environments in which the rotifer could be rescued through the changes in the frequencies of the two algal clonal lines. Additionally, we manipulated the starting frequencies of the two algal clones to investigate the effect of the populations’ defense level on the probability of rescue.

## Material and Methods

### a) Model

We used the model from Becks and colleagues [20] which describes the dynamics of a single predator with two different prey clones in chemostats. The prey clones differ in two functional traits: defense against predation and competitiveness. *C*_*1*_ is the undefended clone with a higher palatability (*p*_*i*_; probability of consumption by the predator) than the defended clone *C*_*2*_. Investing into higher defenses is assumed to lead to a cost of reduced competitiveness, which is modeled here as differences in half-saturation constant (*K*_*C*_), such that *Kc*,_*1*_, of the undefended prey, is smaller than *Kc*,_*2*_, of the defended prey. The predator population, while consisting of one genotype, is divided into two subpopulations, mimicking our experimental system and following previous studies [20,23]. Reproducing predators (*B*) senescent with rate λ and are no longer able to reproduce (*S*) but still consume prey.

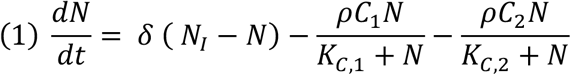

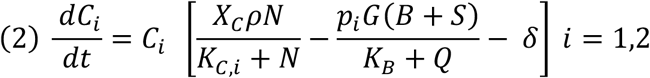

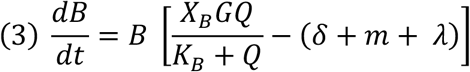

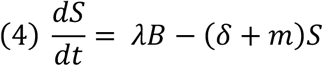

We assume nitrogen (*N*) as limiting resource, which is renewed by a constant inflow with the dilution rate δ and at the concentration *N*_*I*_. Prey population growth and prey nitrogen consumption is modelled as a Monod function with the same nitrogen conversion efficiency *X*_*c*_ and maximum per-capita recruitment *β*_*C*_ for both clones included through the parameter *ρ*=*β*_*C*_ / *X*_*c*_. The nutrient uptake of the prey is modeled to be limited by the half-saturation constants *K*_*C,1*_=2 and *K*_*C,2*_=8. Both prey clones are grazed by predators following a Monod equation which is influenced by their palatability. The higher palatability of the undefended clone (*p*_*1*_=0.8) leads to a higher grazing compared to the defended clone (*p*_*2*_=0.2). The grazing parameter *G*=*β*_*B*_*/ K*_*B*_ is defined by the predator conversion *β*_*B*_ efficiency and predator half saturation constant *K*_*B*_. Predator consumption is limited by the total available prey *Q*=*p*_*1*_*C*_*1*_+*p*_*2*_*C*_*2*_. Reproducing *(B)* and senescent *(S)* predators die at a constant daily mortality rate *m* (all parameters and values see Table S1).

We first identified the steady states values for the densities of predators (*B, S*), and prey clones (*C*_*1*_, *C*_*2*_) with predator background mortality of *m* = 0.055 to use these as initial values for our simulations. To identify these conditions, we ran simulations with combinations of dilution rates (*δ* = 0.1-0.7 by 0.05 d^-1^) and nitrogen concentrations (*N*_*I*_ = 80-400 by 10 μM) as these are the parameter that can experimentally be controlled. As we were interested in scenarios where predator and both prey clones coexist, we only considered those steady states where all three were present with a density over either 0.1 predator/ml and 10^5^ prey cells/ml and when the defended prey had a higher density than the undefended prey.

We started simulations testing for IER using the steady state values for *N, C*_*1*_, *C*_*2*_, *B, S* and varied the starting frequency of the defended clone (0-1, by 0.1) and the mortality rate of the predator (0.055-0.825, by 0.055 d^-1^) in combination with *N*_*I*_ and *δ* (Table S2). For each simulation, we determined whether IER occurred using the following criteria, 1) the predator goes extinct, when the prey clones cannot change in frequency but survives when the prey clones can change in frequency, 2) the minimum predator density is always above 0.1 individuals/ml and 3) the recovery time (i.e., the time needed to reach another maximum after which they established either a new steady state or cycling) was smaller than 100 days. To evaluate the first condition, we calculated the mean defense level of the prey population (frequency *C*_*1*_ x *p*_*1*_ + frequency *C*_*2*_ x *p*_*2*_ from the values of the steady states) and then set *i*=1 in equation (2) with *p*_*1*_ equal to the mean defense and *K*_*C,1*_ equal to the mean half saturation constant. Thus, the prey population could change in density but not their traits. For parameter combinations with longer recovery times than 100 days, IER was considered to be unlikely as the risk of extinction would increase in an experiment and/or with additional environmental noise. An alternative outcome is the extinction of the predator, even when the prey clones can change their frequency or survival in the absence of change in prey frequencies. All simulations were run in R [24] using the R studio environment [25] with the deSolve package [26].

### b) Experiments: cultures

The rotifer population of *Brachionus calyciflorus* used in this study was an obligate asexually reproducing clone, with a generation time of 1.5 to 2 days [20]. The stock was kept by weekly transfers of a small volume to new cultures of *Scenedsmus sp*.. We used two different clonal lines of *Chlamydomonas reinhardtii* as prey. The strains have been isolated from a selection experiment [22] and they differ in their defense against the predator and growth rate. One clonal line has a higher defense against predation and a lower growth rate (CR 1; evolved from strain cc1883 from the Chlamydomonas Resource Center) compared to the other clonal line with a low defense but a higher growth rate (CR 6; evolved from strain cc1144 from the Chlamydomonas Resource Center; Fig. S1). The clones can be distinguished based on their morphology and motility: CR 1 grows in cell aggregates [22] as part of its defense, CR 6 grows mainly as single cells and is motile (Fig. S4). A previous study showed that these traits are heritably over many generations even in the absence of predation [27] and that these traits can generally be linked to specific genetic changes [22]. We refer to predator and prey when referring to the model and defended or undefended algal clone and rotifer for describing the experimental results. To ensure no to little evolutionary change of the algal stock prior to the experiment, they were kept static on agar plates. All cultures were kept at constant light at +/- 20°C in the growth medium [28].

### c) Experiment: test for IER

All IER experiments were started with 4 rotifer/ml and 3.85 * 10^5^ algae cells/ml in tissue cultures flask (TC T-25 standard surface filter cap, Sarstedt) in 21 ml medium. Before adding rotifers to the experimental flask, rotifers from the stock cultures were filtered with a 40 μm cell strainer, left to starve for 3-4 hours and afterwards filtered and washed again to ensure that no *Scenedesmus* was transferred to the experimental cultures. The algal clonal lines were picked two weeks prior the start of the experiment from single colonies from agar plates and grown separately in nitrogen-rich media (800 μM N). Two days before the experiment, they were transferred to medium with 200 μM N to acclimate to the experimental conditions. Rotifers and algae were added together to the culture flasks with medium containing 200 μM N and different NaCl concentration (Table S3). In a previous experiment, we found that rotifers survive NaCl concentration up to 4 mM independent of the algal clone offered, rotifers went extinct at 6 mM NaCl independent of the algae clone, and rotifers survived at 5 mM NaCl when grown with the undefended algal clone and went extinct when grown with the defended algal clone (Fig. S3). Based on these findings, we used 4, 5 and 5.5 mM NaCl for our experiments to test for IER. We mixed the algal clones for different starting frequencies for the defended clone (0, 0.1, 0.3, 0.5, 0.7, 0.9, 1) to test the prediction that the frequency of the defended clone determines the probability of IER. For each combination of NaCl concentration and starting frequency we set up 10 replicates. The experimental populations were kept +/- 20°C in constant light on an orbital shaker (100 rpm, continuous shaking). Each day the inside of the bottles was scratched using an inoculation loop to remove any algae growing on the walls of the flasks. Afterwards, we replaced a third of the culture volume (7 ml) by fresh medium and put the flasks randomly back onto the shaker. From the replaced medium, 1 ml was mixed with Lugol’s iodine to 5% final concentration for algae fixation and 5 ml was used for live rotifer counting. The experiments lasted for eight days, which is approx. four predator generation. We can thus exclude the possibility for evolutionary rescue in the rotifer population due to evolutionary adaptation to increased salinity. Fixated algal samples were counted using a high content microscope (ImageXpress Micro 4®; Molecular Devices) in technical triplicates of 50 μl in 96-well plates with flat bottom with 150 μl of nitrogen free medium. The algae were allowed to sink to the well bottom overnight prior to imaging. We used the CY5 filter with the autofluorescence of the algal cells for imaging, and a Costume Module in MetaXpress® (Molecular Devices) for segmentation and enumeration of algal cells. For acquisition and costume module protocols see SI.

To determine the frequencies of the algal clones, we spread out ∼50 μl of each flask onto two agar plates on day 4 for the 5 mM NaCl and 5.5 mM NaCl, on day 5 for the 4 mM NaCl treatments, as well as on day 8 for all treatments. The plates were then kept at +/- 20°C in constant light to let the algae grow into colonies on the plates. From these, we randomly selected 32 individual colonies and transferred them to 300 μl medium with 800 μM N in 96-well plates and let them grow individually in wells under the same conditions. Each clonal population that grew from a single colony in a well was checked after one week for their morphology by microscopy to identify the clonal population per well (see above).

### d) Experiment: Data and statistical analysis

The theory of IER in a predator-prey system is that the decline in predator population leads to an increase in undefended prey, which then rescues the predator with more available prey. With IER, rotifer density is expected to decline initially before the predator population recovers, creating a u-shaped temporal change in predator density. To take the u-shaped change in population growth into account when determining IER, we estimated rotifer growth rate over the 8 days of the experiment. For which we expected total growth rates close to zero due to the opposing growth rates in IER events. Rotifer growth rates were compared using generalized least of squares (GLS) with NaCl concentration as factorial and the starting frequency of the defended algal clone as numerical predictors. To deal with the heterogeneity in our data we used the varIdent variance structure for the different NaCl concentrations. We compared survival of each rotifer population using GLM with binomial family and with NaCl concentration and the starting frequency of the defended algal clone as predictors. Rotifer populations with a final density below 1 rotifer/ml were considered as extinct. We used survival rates instead of the survival or extinction of each replicate population in Figure 3b.

To detect the shift of in algal frequencies, we used the Sign as the data was neither normally distributed (Shapiro test: W = 0.81624, p-value < 2.2e-16) nor had equal variance (Levene test: Test Statistic = 32.247, p-value < 2.2e-16). Specifically, we compared the differences of the frequencies of the undefended algal clone on day 4 (or 5 for 4mM NaCl) with starting frequencies. The theory of IER assumes that the increase of the undefended prey leads to a positive growth rate of the predator. To test this prediction, we estimated the relationship between the rotifer growth rate of day 4 (or 5) to 8 and the change in density of the undefended algal clone from day 0 to 4 (or 5) using GLS with the same variance structure as above. We tested in the model for an interaction between both predictors to detect any differences in regression depending on the salt concentration. For the 5 mM NaCl data we used a linear regression model (LM) as it was homogeneous. All analysis was performed with base R [24], the BSDA [29], lawstat [30] and the nlme package [31], using the R studio environment [25] and data were plotted with ggplot2 [32].

## 3. Results

### a) Model

The model predicts that predator growth and survival depend on the predator mortality, the possibility for prey evolution and the initial frequency of the defended prey. Predators go extinct with only defended prey independent of the predator mortality (Fig. 1a white area) and survive with only the undefended prey for a large range of predator mortalities (Fig. 1a, blue area). For low predator mortality and initial frequency of the undefended prey ≥ 0.1, the predator and both prey types survive without evolution in the prey (Fig. 1e) but with increasing predator mortality, survival of the predator depends on prey evolution (IER; Fig. 1a purple area, Fig. 1d). The model further predicts that IER becomes more important for predator survival for lower mortality rates when the initial frequency for the defended prey is high (Fig. 1b). Predators go extinct for high mortality independent of the frequency of the defended type. The conditions for IER and rotifer survival on mortality and initial frequency of the defended prey does not change when altering the concentration of the limiting resource and dilution rate (Fig. S5). The probability for predators to survive decreases, however, with increasing nitrogen concentration and dilution rate.

**Figure 1.**
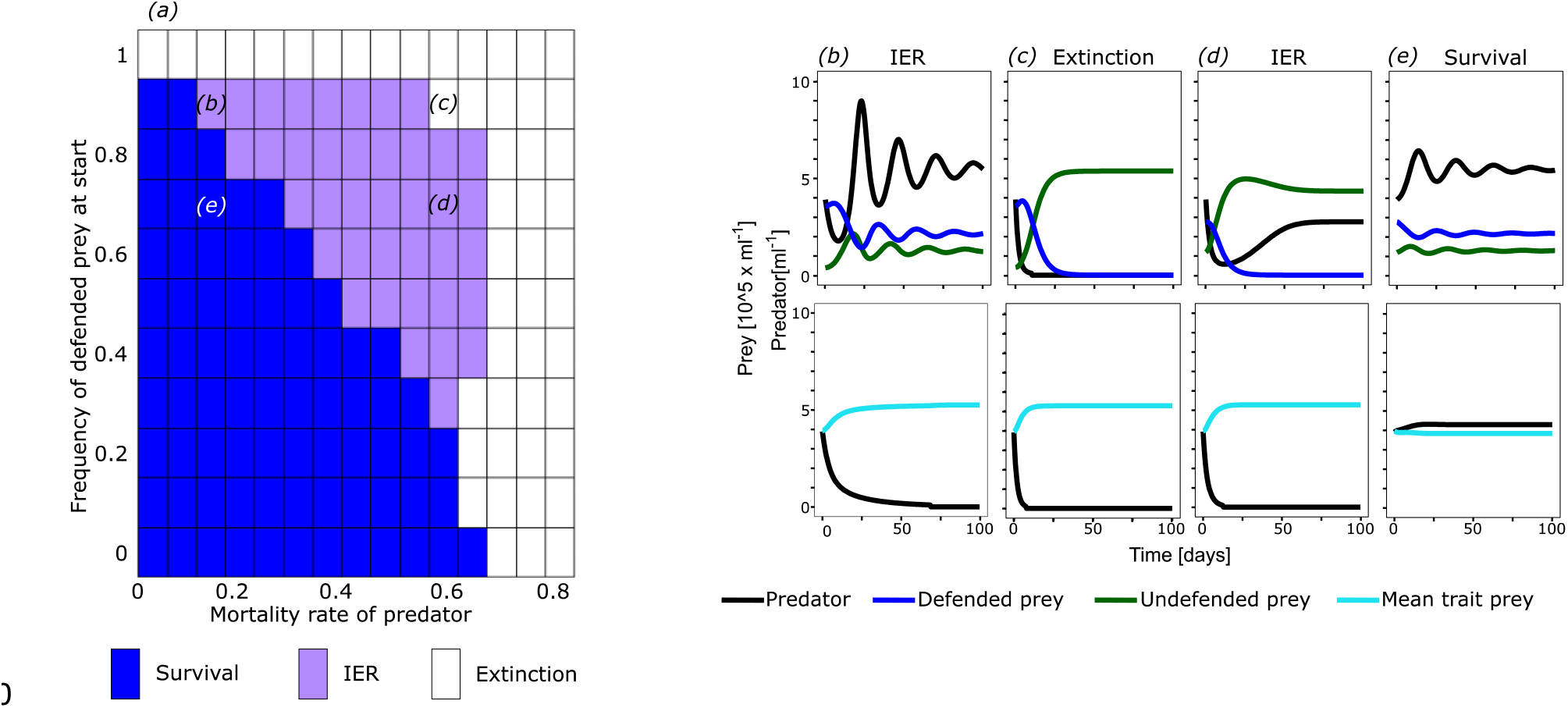
Population dynamics of predator and prey depending on starting frequency of the defended prey clone and mortality rate of the predator from chemostat model. a) Summary of population dynamics: extinction of predator (white), survival of predator without IER (blue) and IER (purple) for simulations with a dilution rate *δ* = 0.3 and nitrogen concentration *N*_*I*_ = 200 μM for a range of mortality rates of the predator and starting frequencies of the defended prey. Panels b-e) show examples of population dynamics for the combinations of predator mortalities and initial frequencies of the defended prey in panel a. Top panels show the simulation result with evolution in the prey population, bottom panels without. Note that the initial defense of the prey population is the same for the evolution (top row) and no-evolution (bottom row) simulations (see Methods). Combinations of mortality rates of the predator and initial frequencies of defended prey for the examples are indicated in a by the corresponding letters.

### b) Experiments

We found that total rotifer growth rate and survival depended on NaCl concentrations and the starting frequency of the defended algal clone (growth rate GLS: NaCl; df: 6, L-ratio: 181,088, p < 0.0001, starting frequency: df: 7, L-ratio: 44.214, p < 0.0001, interaction: df: 9, L-ratio: 41.110, p < 0.001; Fig. 2a; survival GLM: NaCl: df:1, χ^2^ = 83.683, p = 2.2 × 10^−16^, starting frequency: df: 2, χ^2^= 42,058, p = 8.862 × 10^−11^, interaction: df: 2, χ^2^= 25.225, p = 3.33 × 10^−6^ ; Fig. 2b). With NaCl concentrations of 4 mM rotifers population grew independent of the frequencies of the algal clones (Fig. 2a), with only a single extinction event (Fig. 2b). At 5.5 mM NaCl, all populations had negative growth rates leading to low densities or extinction by day 8. The highest survival rates were at low to intermediate starting frequencies of the defended algal clone, but always below 0.5. For 5 mM NaCl, rotifer population growth rates depended on the starting frequencies. At high frequencies of the defended algal clone (≥0.9) rotifer growth rates were all negative, and all but one population went extinct at day 8. At lower frequencies of the defended algal clone, the rotifer population showed no or positive growth, with most populations surviving. Survival rates started declining at the starting frequency of 0.5 for the defended algal clone and no rotifer population survived when growing only with the defended algal clone.

**Figure 2.**
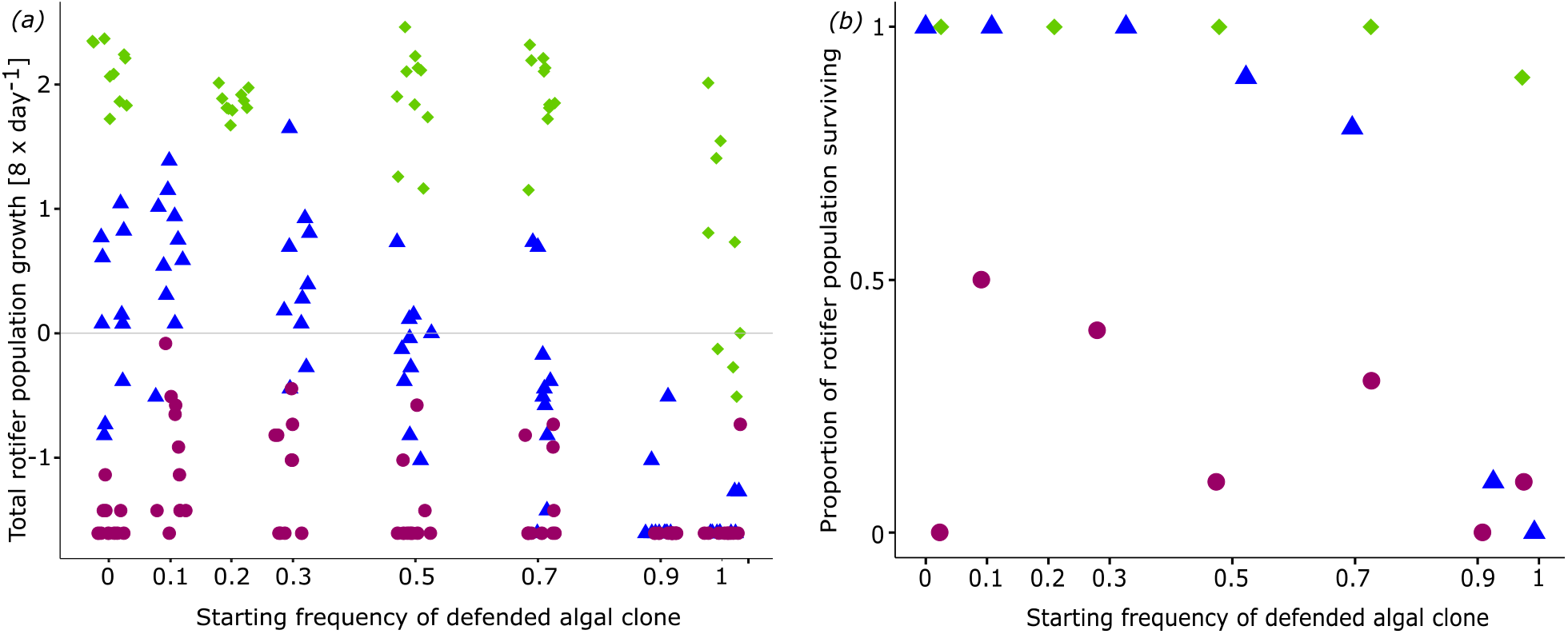
Growth and survival rates of rotifers. (a) Rotifer population growth rates estimated over 8 days when grown in microcosm with different starting frequencies of the defended algal clone and NaCl concentrations (4 mM: green diamond; 5 mM: blue triangle; 5.5 mM: purple circle). Symbols represent replicates (n=10 per starting frequency and NaCl concentration). Grey horizontal line = zero growth rate. (b) Proportion out of 10 rotifer population surviving over the course of the experiment. Values were slightly offset on x-axis for visualization.

Inspecting the temporal dynamics of rotifer and algal clone densities at 5 mM NaCl, we observed the predicted u-shaped dynamics of the rotifers for intermediate initial frequencies of the defended algal clone. For example, at an initial frequency of 0.7 for the defended algal clone (Fig. 3j-s), the density of rotifers decreased during the first days of the experiment, while at the same time the density of the undefended algal clone increased and that of the defended algal clone decreased. This frequency and density change allowed then the rotifer population to recover again with a simultaneous decrease in the undefended algal clone and continuous low density of the defended algal clone (Fig. 3j, m, n, p, q, s). When the algal population consisted only of the defended algal clone, rotifer went extinct after day 6 (Fig. 3a-i) and rotifers survived when treated with only the undefended algal clone (Fig. 3t-dd).

**Figure 3.**
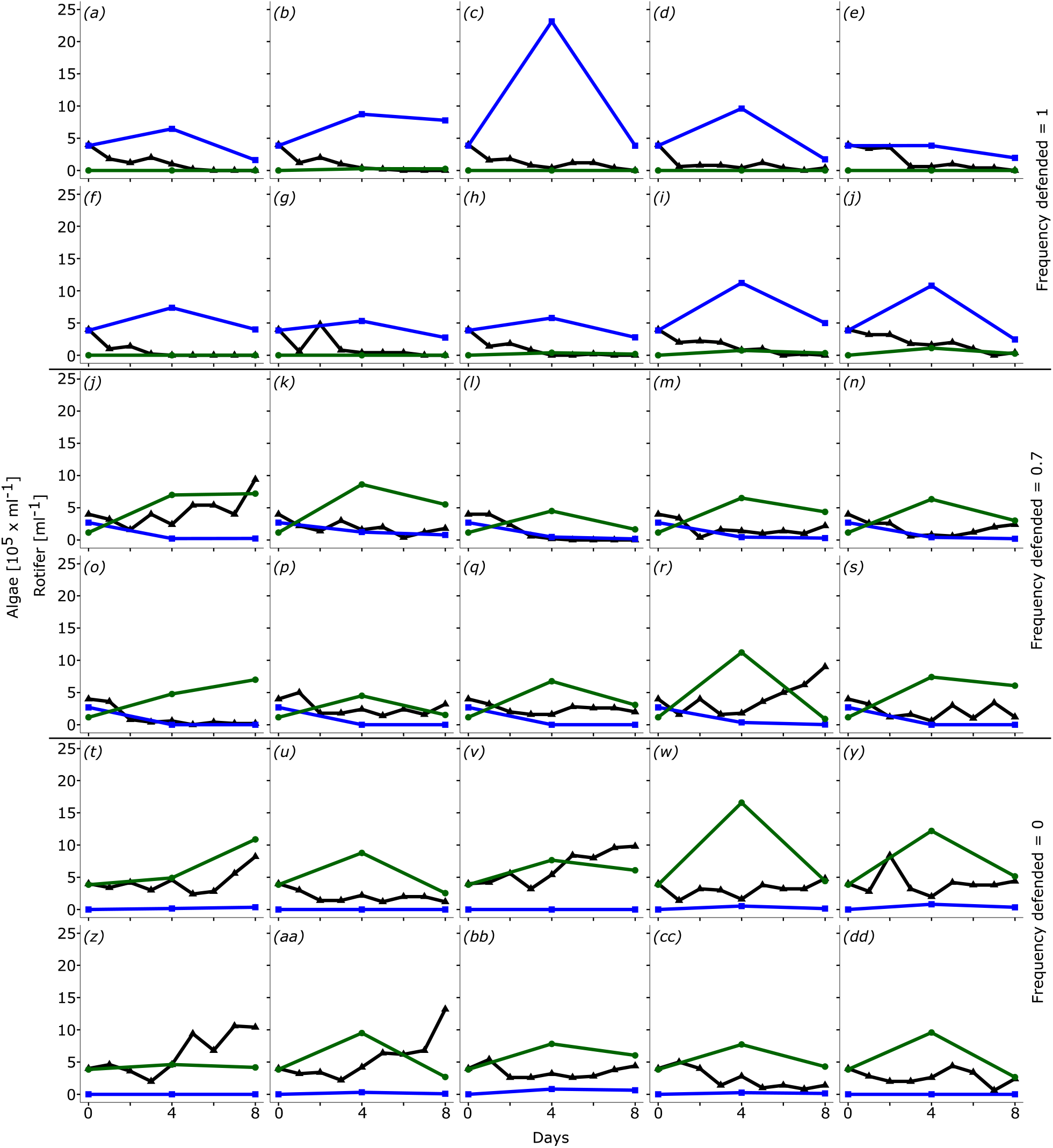
Population dynamics of rotifer and algae population. All replicates shown were grown in 5 mM NaCl with (a-i) starting frequency of the defended algal clone = 1, (j-s) starting frequency of the defended clone = 0.7, (t-dd) starting frequency of the defended clone = 0. Density of rotifers (black triangle), the defended algal prey (blue square) and undefended algal prey (green circle).

The frequencies of the algal clones shifted at the first half of the experiment when both algal clones were present (Sign test, S = 23, p < 2.2 × 10^−16^). To test whether this shift contributed to the predator rescue, we estimated the relationship between the changes in rotifer growth rate from day 4 (or 5) to 8 with earlier changes in the density of the undefended algal clone from day 0 to day 4 (or 5). We used density rather than frequency as the rotifer growth depends on food quality and quantity. We found that the increase in rotifer growth rate followed the preceding increase in the density of the undefended algal clone (GLS: NaCl effect: df: 6, L-ratio: 13.564, p = 0.001, change in undefended algal density: df:7, L-ratio: 11.665, p = 6 × 10^−4^; interaction: df: 9, L-ratio: 10.437, p: 0.0054; Fig. 4). Examining the dependencies for the different salt concentrations, we found a positive regression line at 5mM NaCl but not for the other NaCl concentrations.

**Figure 4.**
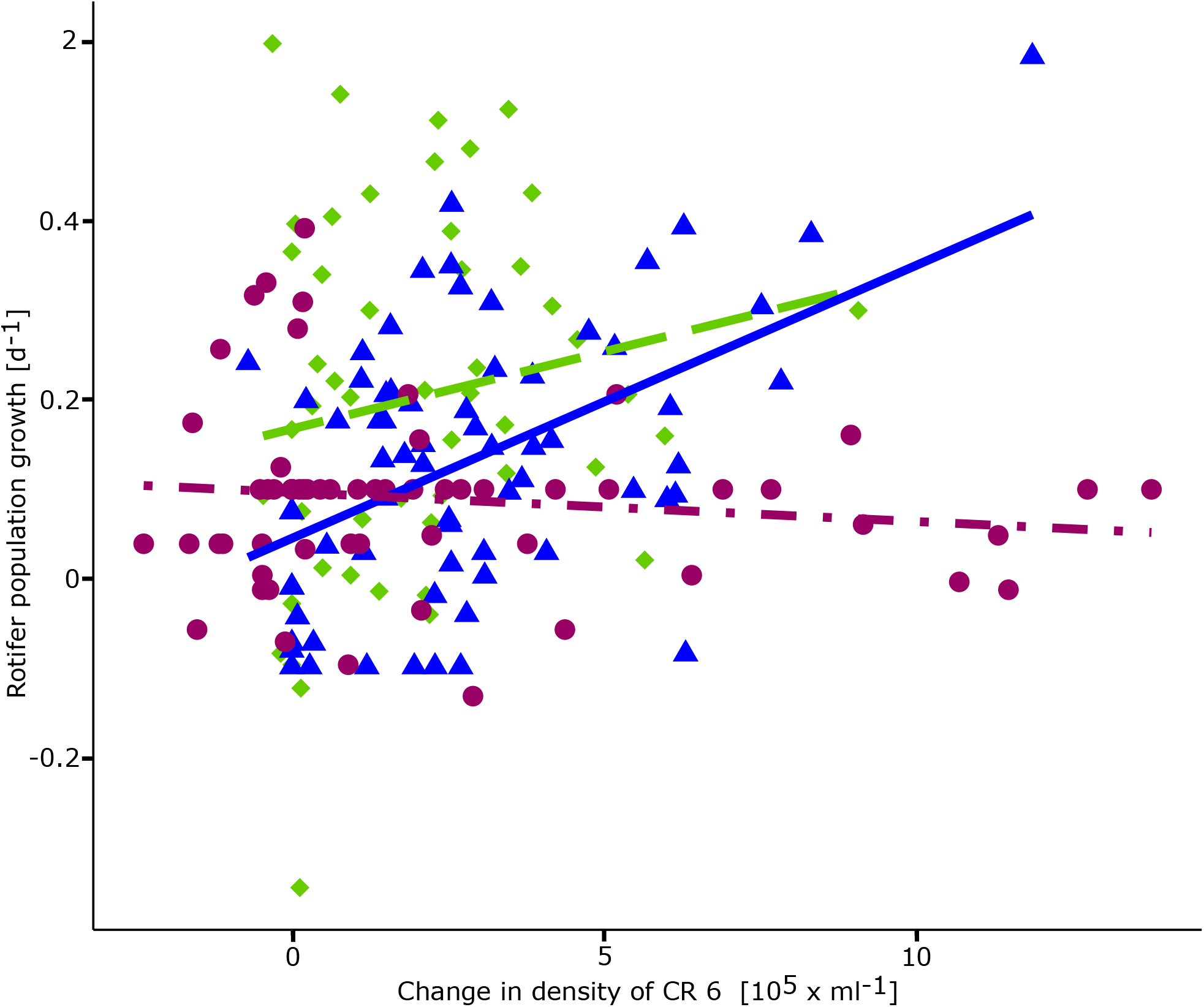
Rotifer growth depending on preceding change in density of undefended algae. Regression lines between rotifer population daily growth rate (from day 4 or 5 to 8 of the experiment) and changes in density of the undefended algae (estimated between day 0 and 4 or 5) for different NaCl concentrations (4 mM: green diamond and green dashed line; 5 mM: blue triangle and blue solid line; 5.5 mM: purple circle and purple dot-dashed line; including all treatments of the different starting frequencies). Symbols represent replicates (n=10 per starting frequency and NaCl concentration).

## 4. Discussion

IER is a process that allows a population to persist through evolution in an interacting species in environments where they would otherwise go extinct [12]. We tested IER in model simulations and experiments with predator and its prey with the goal to determine conditions under which IER can facilitate survival of the predator. Consistent with our predictions, we found that IER is possible with increased predator mortality and that the frequency with which IER can be observed was depending on the initial frequency of defended prey.

Our results show similarities to ER with respect to observing rescue as a function of how the absolute fitness of the population changes with the increase of a genotypes’ frequency [33,34]. In ER the genotype increases due to its increased fitness in the new environment [1–3] while in IER, the abundance of the undefended clone increases when predator density declines as it is the more competitive clone. Thus, the general conditions favoring ER [33] should also favor IER. Specifically, we explored here the role of the initial frequency (i.e., standing genetic variation) of the undefended prey and found in model and experiments, that predator rescue did not occur, when the initial frequency of the undefended prey was too low. Low starting frequency of the undefended algal clone delayed its increase so that the rotifer population went extinct before the density of the undefended algal clone increased enough to avert extinction of the rotifer population. At higher starting frequencies of the undefended algal clone, the increase of the undefended algal clone was sufficiently fast to rescue the rotifer population or the undefended algal clone densities were sufficient from start to sustain the rotifer population despite the increased mortality, leading to positive rotifer growth.

We did not test for the time to minimum population size, which is a function of the initial prey frequency as the predator extinction was very fast in our experiments. IER and the time to minimum population size is also expected to be a function of the fitness advantage of the rescue genotype in the new environment. While we only tested IER for one combination of defended and undefended prey (algal clone), IER should depend on trait combinations of the prey clones and the cost of defense. A cheap defense, where high levels of defense correlate with a small reduction in growth rate, should increase time to minimum population size, thus decreasing the chance for IER. While a costly defense should decrease the time to minimum population size, which would increase the probability of observing IER. However, when long-term predator-prey dynamics are considered, the persistence of prey diversity depends on the cost of defense [27,35], thereby allowing for the possibility of IER.

We observed a pattern resembling a selective sweep in the algal population, where the undefended clones’ frequency increased to almost 1 during IER at high mortality rates. While the reduction of phenotypic diversity in the prey population allows the persistence of the predator, it can also alter the population dynamics of the system [20,23,36] rendering it more susceptible for future perturbations. Another outcome of IER was that defended and undefended prey coexisted, but that the frequency of the defended prey decreased compared to the cases where there was no additional predator mortality and IER. In the latter cases, the predator mediated maintenance of diversity in the prey population was still present [10,21]. As trait diversity can contribute to stabilizing ecosystem functioning [14,37], understanding the conditions that determine the outcome of IER will help to better predict responses to recurrent or consecutive environmental changes.

Changes in dilution rate and nitrogen concentration did not affect whether IER occurs in our model (Fig. S5). We only examined here a small fraction of the parameter space with combinations for dilution rate and nitrogen concentrations for which we found steady state conditions without additional mortality. This allowed us testing the model predictions in short-term experiments without long transient periods. Whether the probability for observing IER changes when the system is cyclic requires additional studies but a previous study showed that the robustness of the population dynamics of a predator-prey system in the presence of a continuous disturbance is strongly impacted by the speed of adaptation in prey and/or predator population [38]. Furthermore, cyclic population dynamics in a predator-prey system with evolution in the prey population exhibit typically anti-phase cycles [23,38]. Under these conditions, the defended prey dominates the prey population when predator densities are at their peak, and the undefended prey dominates when predator densities are lowest. IER should be possible when the environment changes during high densities of the predator as the decline of the predator will lead to the increase of the undefended prey. If the environment changes during low densities of the predator, it should sustain due to the high density of the undefended prey.

Phenotypic plastic responses in the rotifer and algal population may also contribute to rescue of the predator [12]. We consider this possibility for inducible changes in the rotifer and algal population as negligible because the rotifer populations showed no growth changes in their controls with only one clone present, e.g. they grew to high density with only the undefended algal clone at 4 mM NaCl and went extinct with only the defended algal clone at 5 and 5.5 mM NaCl. Furthermore, we previously showed the heritability of the traits in the algal clones and linked these to genetic changes [22] and that the morphology (here used to distinguish the two prey clones as well as determining their interaction and with that of the rotifer) was stable at least for nine generations [27], which covers the timescale of our experiment.

Understanding how populations respond to environmental changes is essential for predicting their ecological and evolutionary dynamics as well as their persistence. Evolutionary responses that rescue populations from extinction after environmental changes can be important driver for the maintenance of ecosystems processes and functions [39], but research on ER is still mostly focused on single species [40]. Only few studies on ER have incorporated species interactions and community dynamics and it is currently unclear which conditions influence the probability of ER. For example, studies on mutualistic interactions [16] have shown a decrease in the probability of ER while studies focusing on antagonistic interactions [14,15,17] found a decrease or increase in the probability of ER. An important determinant for ER in communities is how many species are threatened by extinction and their interactions within the community. Mismatches between responses of for example consumer and their resources can lead to the extinction of species that are not directly affected or to a lesser amount by the environmental change, when their resource declines [41–43]. Incorporating evolutionary responses in one or more species could impact the probability survival of a threatened population [44], either through the mechanism of IER as studied her, through effects on coevolution [11,44,45] or changes in selection and demographic changes [46].

## 5. Authors’ Contribution

R. H and L.B. designed the experiment. R.H. ran the experiment, analyzed the data, and wrote the first draft of the manuscript. R.H. and L.B. wrote the final version of the manuscript.

## 8. Acknowledgments

We thank Eva Lievens for help with the data analysis, Tom Reveillon for help with the method development and Natascha Handke and Tamara Wonner for help with the experiment.

## Supplemental Material

**Table S1.** Parameters for predator-prey model

| Parameter | Description | Value |
| --- | --- | --- |
| $N_I$ | Limiting resource | 80-370 $\mu\text{M}$ |
| $\delta$ | Dilution rate | 0.1-0.7 / d |
| $m$ | Predator mortality | 0.055-0.825 |
| $\lambda$ | Predator senescence rate | 0.4 / d |
| $X_C$ | Prey conversion ($10^9$ cells/$\mu\text{mol N}$) | 0.0027 |
| $\beta_C$ | Maximum prey per-capita recruitment | 0.72 / d |
| $\rho$ | $\beta_C / X_C$ | 270 $\mu\text{M N}/10^9$ cells * d |
| $\beta_B$ | Maximum predator per-capita recruitment | 1.9 / d |
| $X_B$ | Predator conversion | 170 predator / $10^6$ prey cells |
| $K_B$ | Predator half-saturation constant | $0.15 \times 10^6$ prey cells / ml |
| $G$ | Predator grazing rate parameter = $\beta_B / K_B$ | 0.011 ml / predator * d |

**Table S2.** Nitrogen concentrations (μM; N), prey densities (10^6^ cells/ml; C1, C_2_) and predator densities (Ind/ml; B, S) for each steady state with the corresponding nitrogen concentration in the reservoir N_I_ and daily dilution rate δ.

| N | C<sub>1</sub> | C<sub>2</sub> | B | S | N<sub>i</sub> | $\delta$ |
| --- | --- | --- | --- | --- | --- | --- |
| 8.26 | 0.10 | 0.11 | 3.83 | 4.32 | 130 | 0.3 |
| 3.67 | 0.03 | 0.23 | 2.60 | 5.22 | 170 | 0.15 |
| 3.68 | 0.02 | 0.26 | 2.72 | 5.33 | 180 | 0.15 |
| 4.93 | 0.03 | 0.28 | 3.30 | 5.09 | 180 | 0.2 |
| 3.46 | 0.01 | 0.32 | 2.54 | 4.86 | 190 | 0.15 |
| 4.99 | 0.02 | 0.32 | 3.28 | 5.14 | 190 | 0.2 |
| 6.56 | 0.03 | 0.32 | 3.69 | 4.85 | 190 | 0.25 |
| 4.99 | 0.01 | 0.35 | 3.27 | 5.13 | 200 | 0.2 |
| 6.49 | 0.02 | 0.36 | 3.64 | 4.76 | 200 | 0.25 |
| 8.41 | 0.04 | 0.36 | 3.90 | 4.42 | 200 | 0.3 |
| 4.86 | 0.00 | 0.40 | 3.10 | 4.86 | 210 | 0.2 |
| 6.50 | 0.01 | 0.40 | 3.64 | 4.77 | 210 | 0.25 |
| 8.32 | 0.03 | 0.39 | 3.87 | 4.35 | 210 | 0.3 |
| 6.50 | 0.01 | 0.43 | 3.64 | 4.77 | 220 | 0.25 |
| 8.34 | 0.02 | 0.43 | 3.87 | 4.36 | 220 | 0.3 |
| 10.59 | 0.04 | 0.42 | 3.98 | 3.91 | 220 | 0.35 |
| 8.34 | 0.01 | 0.47 | 3.87 | 4.36 | 230 | 0.3 |
| 10.63 | 0.03 | 0.46 | 3.96 | 3.92 | 230 | 0.35 |
| 8.37 | 0.00 | 0.50 | 3.91 | 4.41 | 240 | 0.3 |
| 10.62 | 0.02 | 0.49 | 3.97 | 3.92 | 240 | 0.35 |
| 13.47 | 0.04 | 0.48 | 3.86 | 3.39 | 240 | 0.4 |
| 10.63 | 0.01 | 0.53 | 3.97 | 3.92 | 250 | 0.35 |
| 13.65 | 0.03 | 0.52 | 3.97 | 3.49 | 250 | 0.4 |
| 10.72 | 0.00 | 0.56 | 4.06 | 4.01 | 260 | 0.35 |

Indirect evolutionary rescue
| N | C1 | C2 | B | S | N<sub>i</sub> | δ |
| --- | --- | --- | --- | --- | --- | --- |
| 13.57 | 0.02 | 0.55 | 3.92 | 3.45 | 260 | 0.4 |
| 13.58 | 0.01 | 0.59 | 3.93 | 3.46 | 270 | 0.4 |
| 13.60 | 0.00 | 0.62 | 3.95 | 3.47 | 280 | 0.4 |
| 17.48 | 0.03 | 0.60 | 3.73 | 2.95 | 280 | 0.45 |
| 17.63 | 0.02 | 0.64 | 3.79 | 3.00 | 290 | 0.45 |
| 17.59 | 0.01 | 0.68 | 3.77 | 2.98 | 300 | 0.45 |
| 23.12 | 0.01 | 0.76 | 3.39 | 2.44 | 330 | 0.5 |
| 24.00 | 0.00 | 0.78 | 3.78 | 2.72 | 340 | 0.5 |
| 31.88 | 0.02 | 0.82 | 2.97 | 1.95 | 360 | 0.55 |
| 31.91 | 0.00 | 0.86 | 2.91 | 1.93 | 370 | 0.55 |

**Table S3.** Experimental conditions.

| NaCl<br>[mM] | μM N | Starting Frequency of<br>CR 1 | Rotifer /<br>ml | Algae cells / ml |
| --- | --- | --- | --- | --- |
| 4 | 200 | 0, 0.2, 0.5, 0.7, 1 | 4 | 3.85 x 10<sup>5</sup> |
| 5 | 200 | 0, 0.1, 0.3, 0.5, 0.7, 0.9, 1 | 4 | 3.85 x 10<sup>5</sup> |
| 5.5 | 200 | 0, 0.1, 0.3, 0.5, 0.7, 0.9, 1 | 4 | 3.85 x 10<sup>5</sup> |

**Figure S1.**
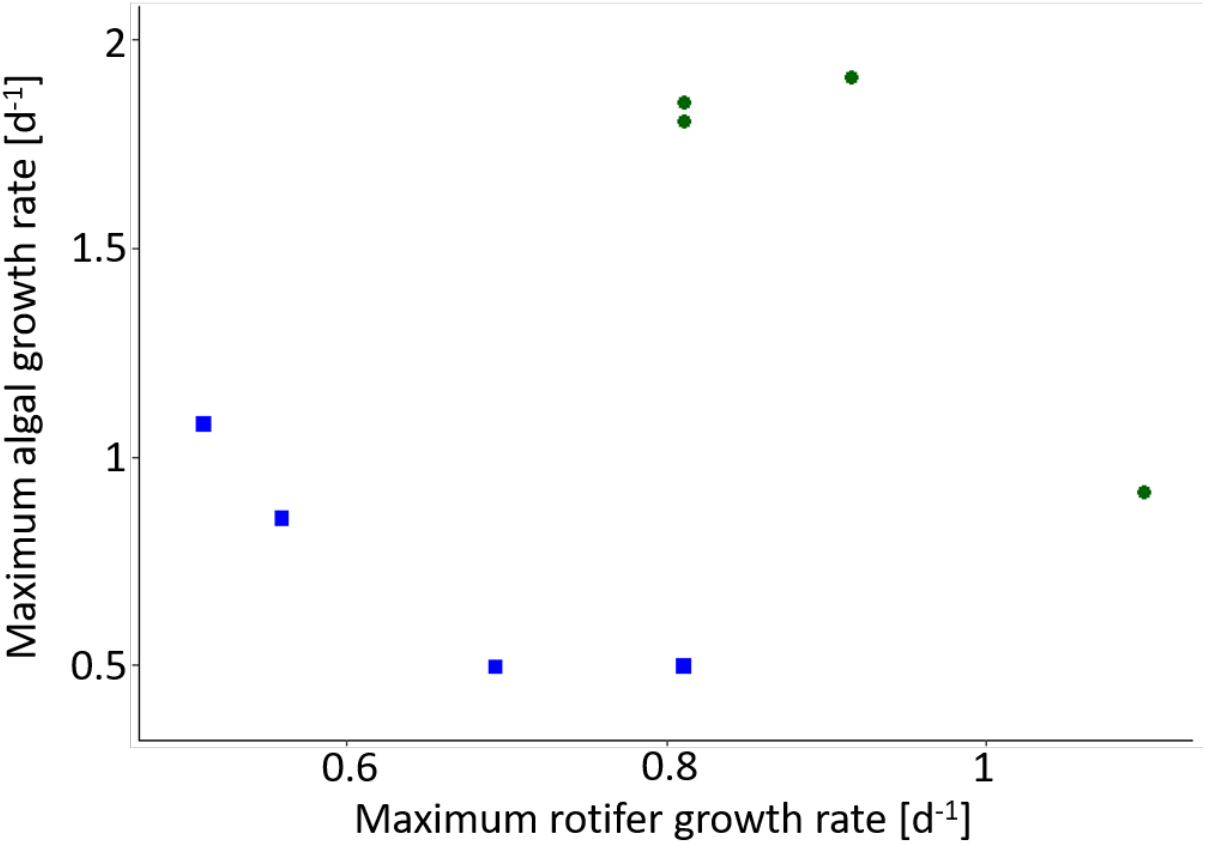
Correlation between algal growth rate and defense against predation (measured by rotifer growth rate in presence of one clone) of CR 1 (blue square) and CR 6 (green circle). Each symbol represents one replicate. The maximum daily growth rate of the algal clones was taken from an eight-day long growth experiment and the maximum rotifer growth rate was taken from a five-day long growth experiment.

**Figure S2.**
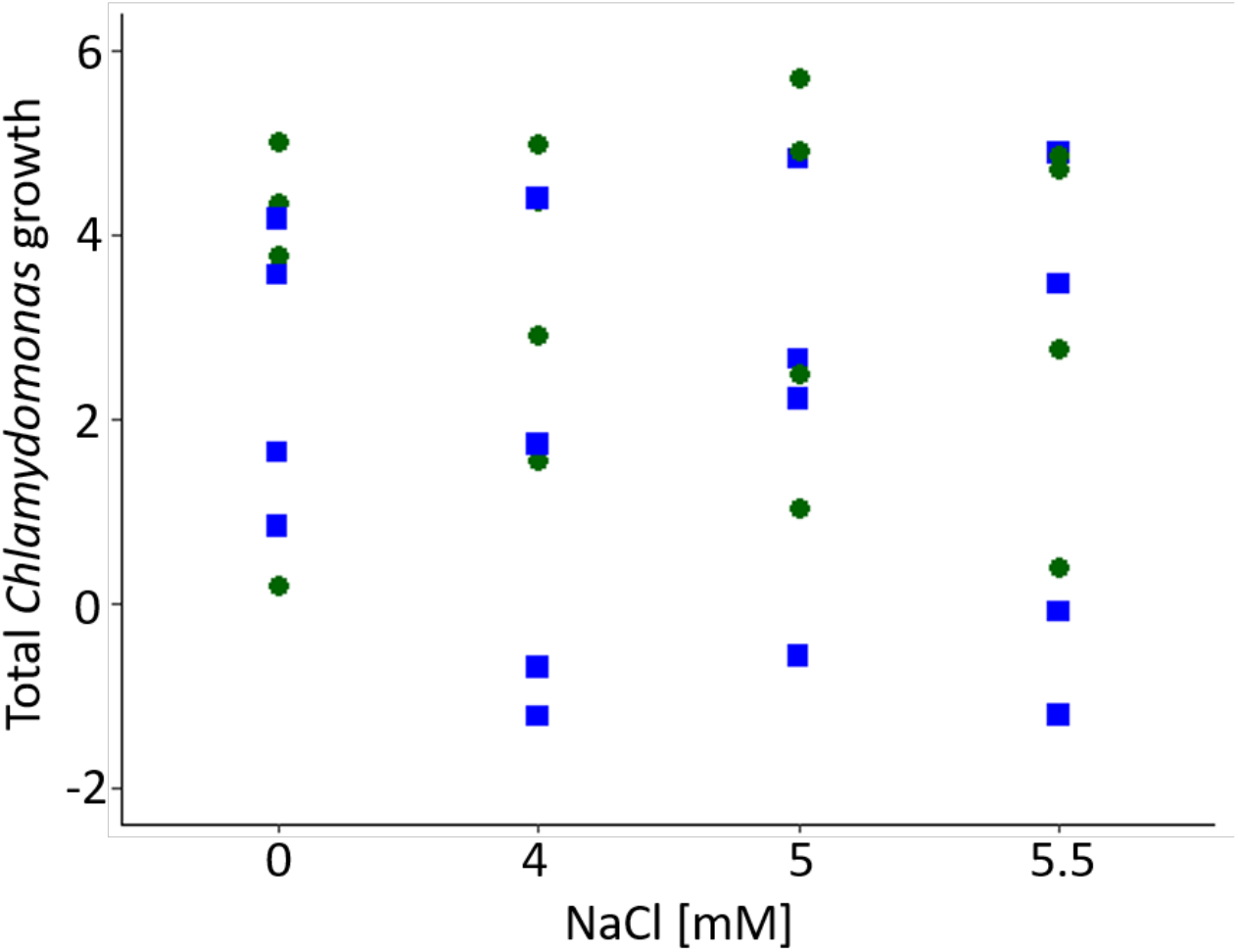
Growth rate over four days of CR 1 (blue squares) and CR 6 (green circles). Each symbol represents one replicate. No significance was found for the effect of NaCl in the growth rate of the clones (ANOVA with NaCl concentration as explanatory variable: F_1,29_=0.1469, p=0.70; *n*=4).

**Figure S3.**
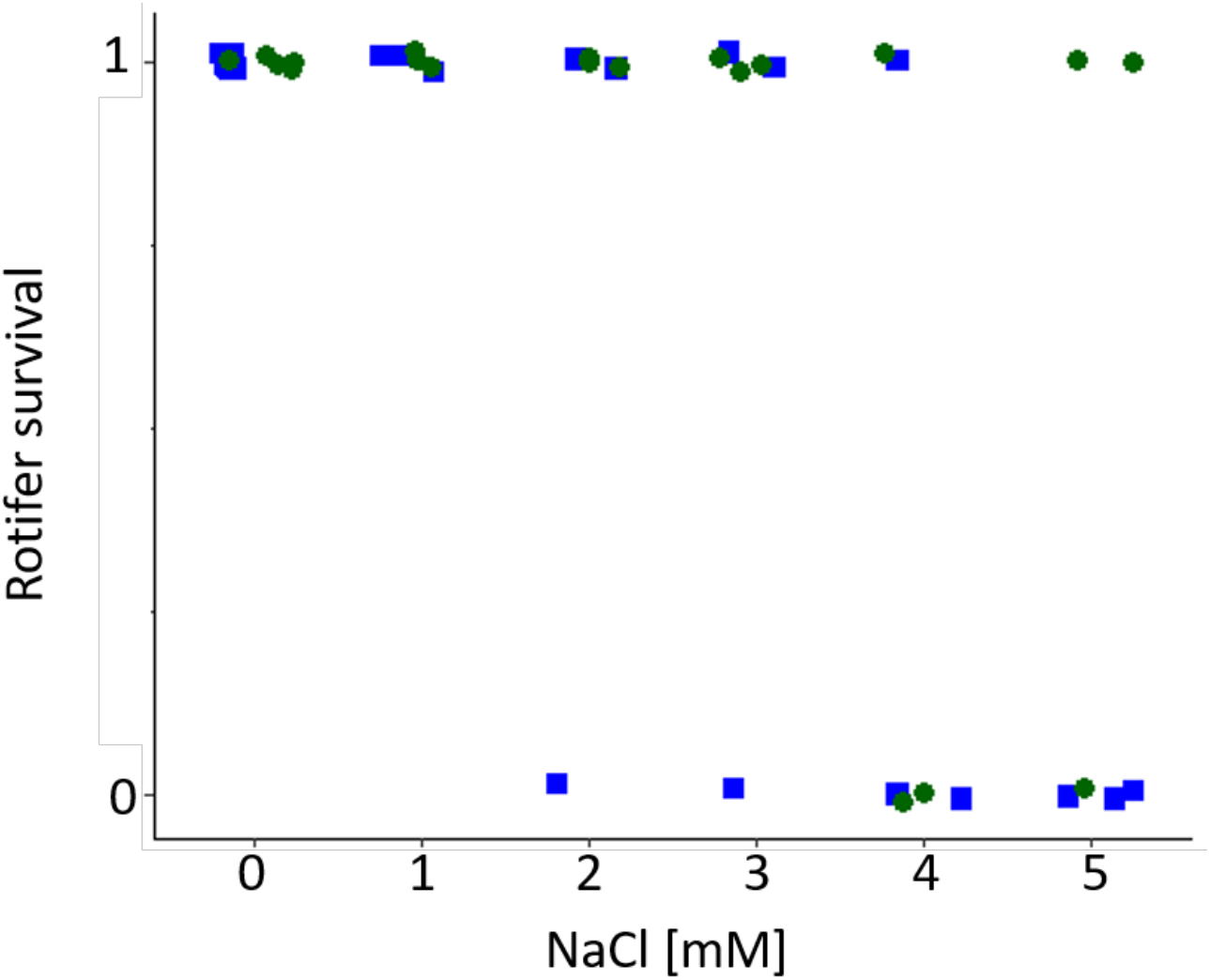
Rotifer survival when grown for six days with CR 1 (blue square) or CR 6 (green circle) (*n* = 3). Each symbol represents one replicate. Rotifers were considered as extinct if the density was below 1 rotifer/ml.

**Figure S4.**
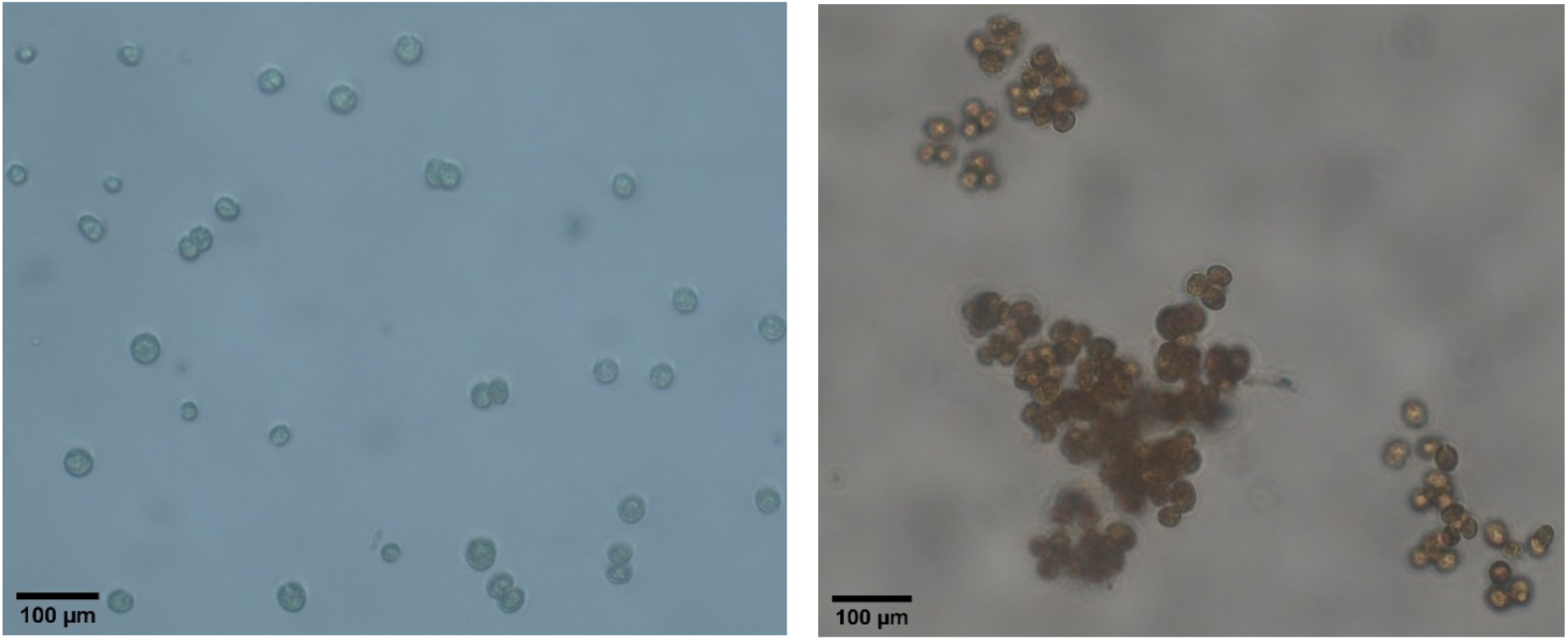
Representative morphotypes of the algal clones CR 1 and CR 6. CR 1 (left) forms bigger clumps while CR 6 (right) is typically single celled. Algal samples were fixated with Lugol’s iodine solution prior to imaging. Scale bar implemented with ImageJ.

**Figure S5.**
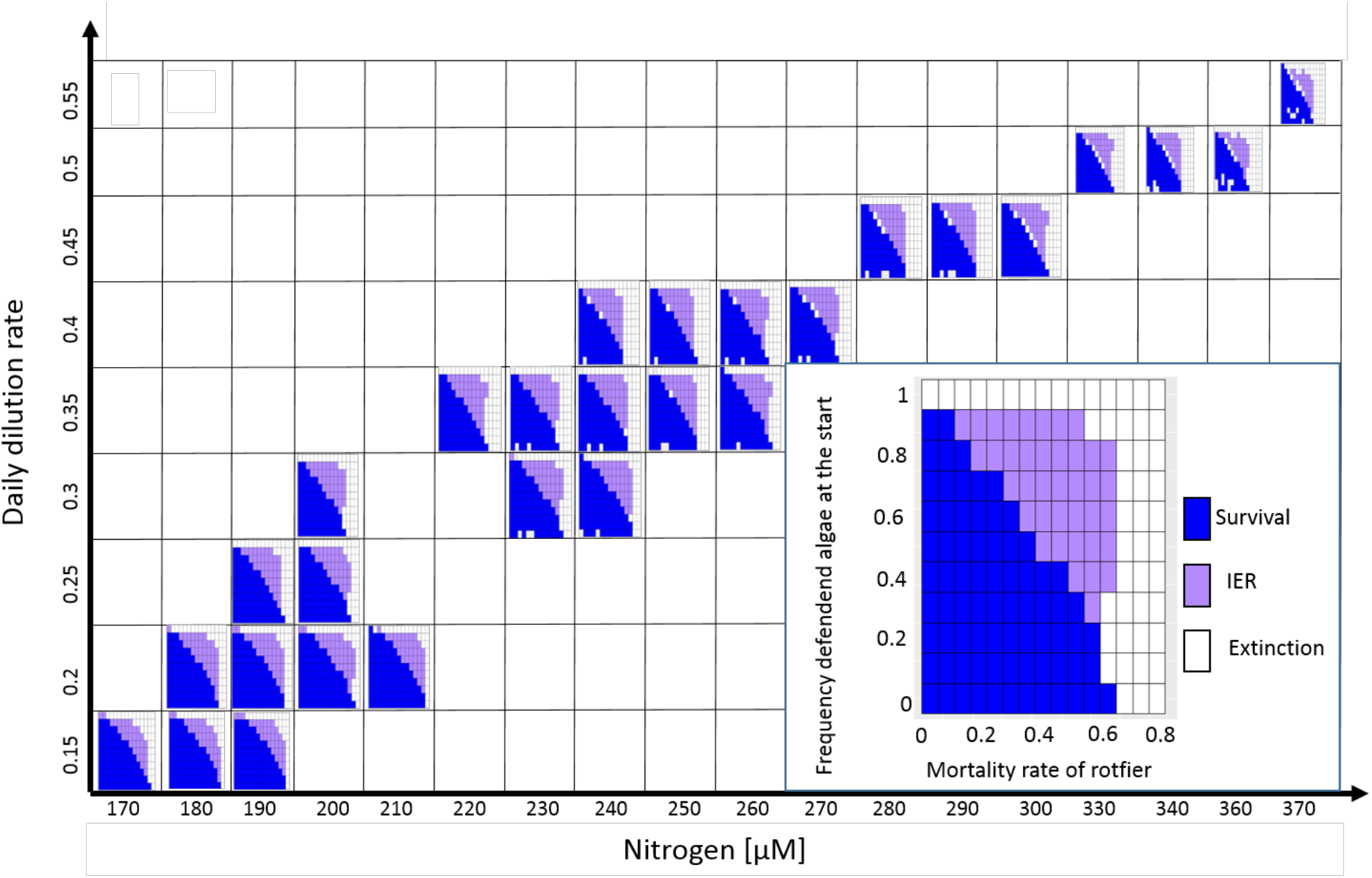
Combinations of N_l_ and ***δ*** of the steady states (Table S2) where determined over a range of starting frequencies of the defended prey clone and mortality rate of the predator. Outcomes of the simulations were either survival of the predator without prey evolution (blue), IER (purple) or extinction of the predator with evolution of the prey (white). In the right bottom corner the results from N_I_ = 200 and ***δ*** = 0.3 are shown for clearance. Other parameters were used according to Table S1. Some steady states had to be excluded as the modeling process led to errors.

